# The inceptionist’s guide to base editing – *de novo* PAM generation to reach initially inaccessible target-sites

**DOI:** 10.1101/2022.07.07.499158

**Authors:** Kaisa Pakari, Joachim Wittbrodt, Thomas Thumberger

## Abstract

Base editing by CRISPR crucially depends on the presence of a PAM in proper distance to the editing-site. Here we present and validate an efficient one-shot approach termed “inception”, relaxing these constraints. This is achieved by sequential, combinatorial base editing: *de novo* generated synonymous, non-synonymous or intronic PAM-sites facilitate subsequent base editing at nucleotide positions inaccessible before, increasing the targeting range of highly precise editing approaches.

## Introduction

The highly precise base editing becomes progressively more important for therapeutic applications as apparent by the first clinical trials launched (Eisenstein, 2022). Its general applicability is constrained by the intrinsic geometry of the target-site: a protospacer adjacent motif (PAM) sequence has to be located in the correct distance, i.e. 13-17 nucleotides (nt) downstream to the desired nucleotide target for proper base editing. One attempt to overcome this limitation currently considered is the use of so-called (near) PAM-less (Walton et al., 2020) or PAM-free (Tan et al., 2022) base editors. However, their extended targeting range comes at cost of reduced specificity (Walton et al., 2020).

Here we present an alternative, taking advantage of the well-established adenine and cytosine base editors (ABE8e (Richter et al., 2020), ancBE4max (Koblan et al., 2018), evoBE4max (Thuronyi et al., 2019)) with NGG-PAM recognition to reach previously inaccessible base editing sites. This is achieved by an intermediate *de novo* PAM generation and subsequent upstream base editing that we termed “inception”.

## Results and Discussion

Based on a canonical guide RNA target-site, a *de novo* PAM can be introduced by A-to-G base editing if the adenosine(s) of AA, GA or AG dinucleotides are contained within the canonical base editing window. Conditioned by this editing, a novel guide RNA target-site becomes available for a further guide RNA/base editor complex, introducing the intended mutation 27-36 nucleotides upstream of the canonical PAM (Fig. 1A). In other words, for an edit of interest a canonical NGG PAM can be located anywhere within a distance of 27-36 nucleotides downstream. This flexibility relaxes the intrinsic constraints of a single target site while keeping the targeting specificity. The sequential editing nature of the inception approach allows to apply all at once: the A-to-G and a possible additional base editor as well as the canonical and inception guide RNAs. Comparing the top ten most studied genes in human and their orthologs in commonly used model organisms, inception increases the number of editing sites by 65 % (Fig. 1B, Table S1).

**Figure 1.**
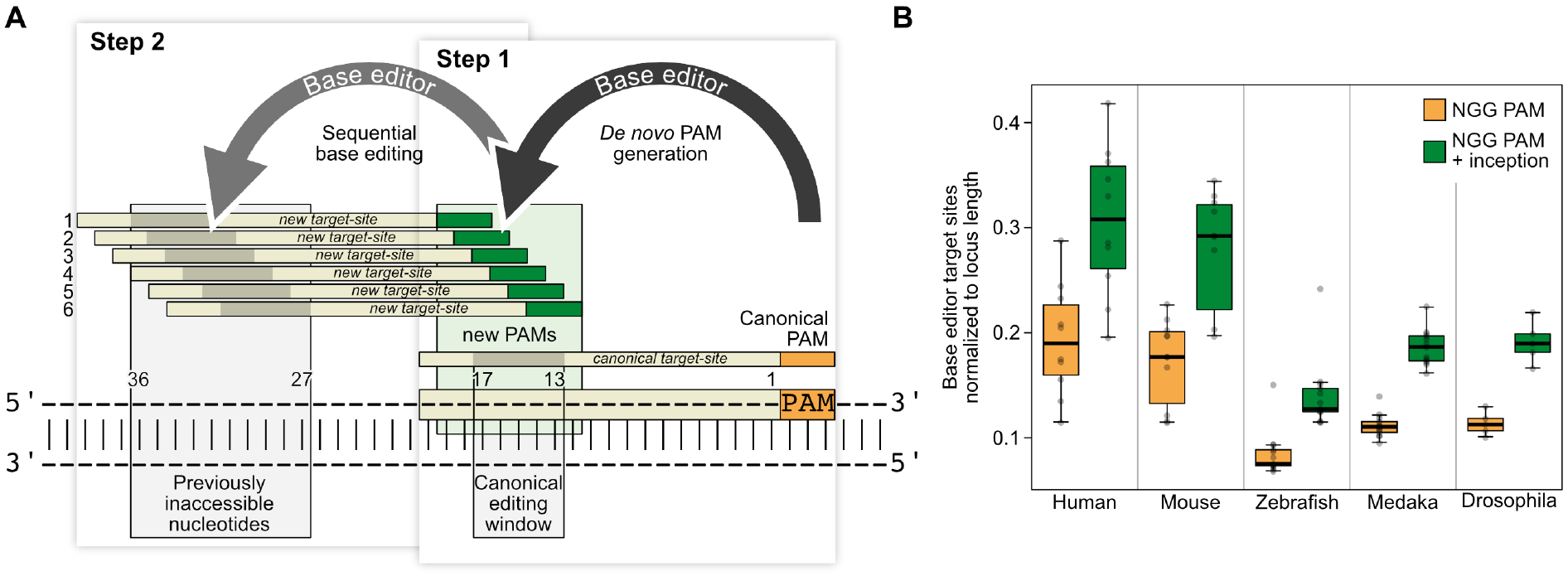
Inception increases the base editing scope by *de novo* PAM generation. (A) Schematic representation of the one-shot inception approach (simultaneous application of two base editors and two guide RNAs). If the adenosine(s) of NGA, NAA and NAG motifs are contained within the canonical base editing window, an A-to-G edit event leads to the generation of (up to six) new PAM(s) (green, Step 1), rendering a new guide RNA target-site available for a second base editing event (Step 2) 27-36 nucleotides upstream of the canonical PAM (orange). As the second base editing relies on the first, the inception approach depicts a locally confined sequential editing event. (B) Abundance analysis of canonical base editor target sites with NGG PAM (orange) and target-site increase upon inception approach (green) normalized to gene locus length. Base editor target sites contain A or C nucleotide(s) in the respective editing window. Comparison of the top ten studied genes in human (*AKT1, APOE, EGFR, ESR1, IL6, MTHFR, TGFBI, TNF, TP53, VEGFA*) and orthologs of commonly used model organisms (Table S1). Across all loci and organisms, inception increases accessible target-sites by 64.8 % ± 6.0 %. PAM, protospacer adjacent motif

To investigate a general applicability of the inception concept, we applied this serial targeting approach in three different settings: two knock-out regimes by introducing a pre-termination STOP codon (PTC) in an open reading frame or the removal of a splice-acceptor-site by targeting intronic sequences. Third, we used inception to introduce locally confined, predictable multi-codon changes to generate allelic variances with different phenotypic severity.

To address the efficiency of a knock-out via inception, we targeted the well described *oculocutaneous albinism 2* (*oca2*) gene responsible for the pigmentation of the retinal pigmented epithelium (RPE) in the Japanese rice fish medaka (*Oryzias latipes*) (Cornean et al., 2022, Lischik et al., 2019). The loss of pigmentation depends on bi-allelic editing of *oca2* which we use as proxy to determine the knock-out efficiency via an established analysis pipeline (Thumberger et al., 2022). While the *de novo* PAM generation depends on A-to-G substitution, the subsequent PTC is introduced by C-to-T editing (Fig. 2A). Guide RNA target-sites were chosen such that the canonical site is expected to generate synonymous edits, and the subsequent targeting to introduce the PTC.

**Figure 2.**
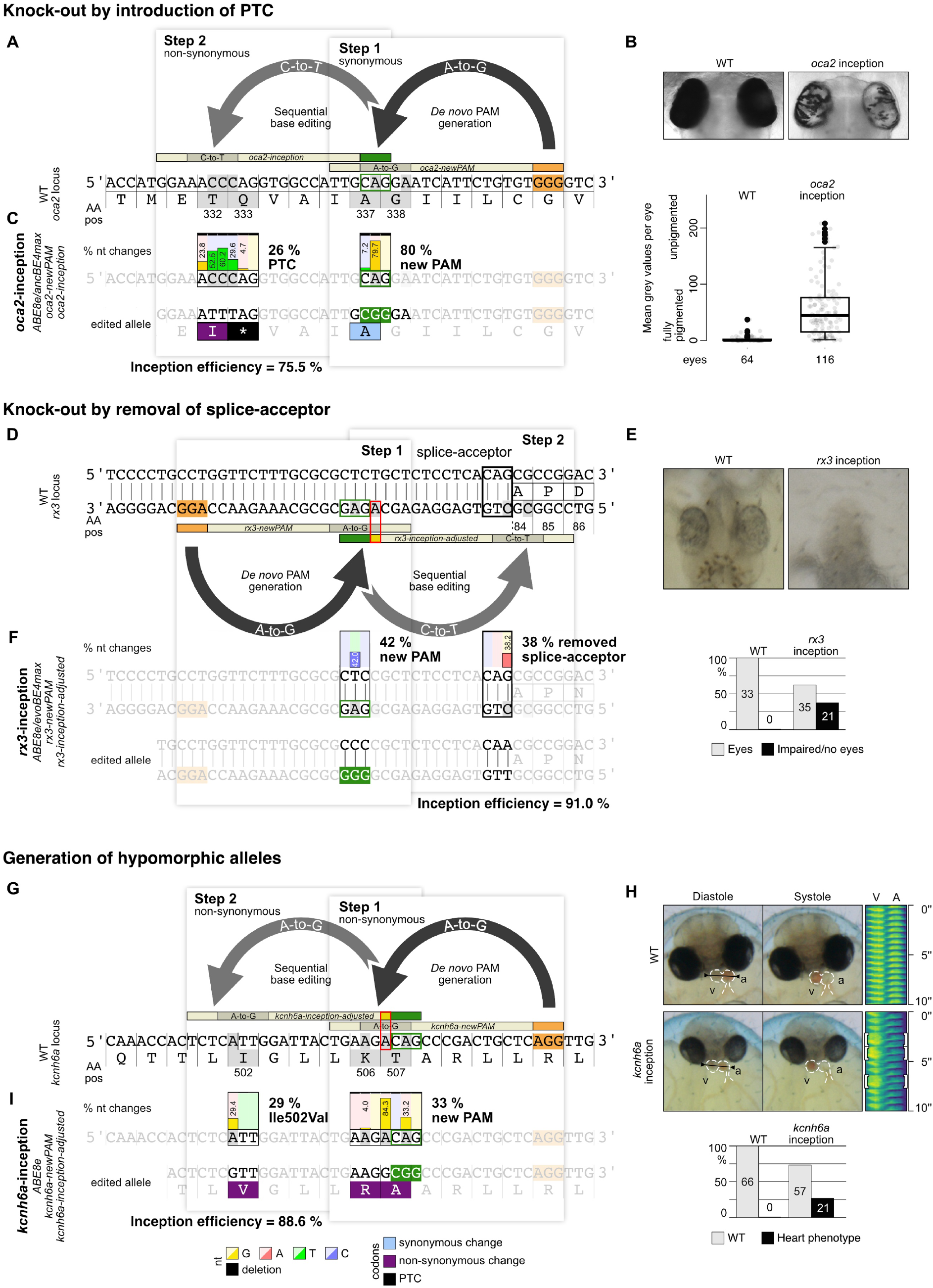
Inception approaches efficiently introduce knock-out mutations and locally confined multi-codon changes. (A) Introduction of a pre-termination STOP codon (PTC) via inception at *oca2* p.Q333 by synonymous *de novo* PAM generation (ABE8e, CAG>CGG, p.Ala337Ala) and subsequent non-synonymous editing (ancBE4max, CAG>TAG, p.Q333*). (B) At 4.5 days post fertilization (dpf), full pigmentation of wild-type eyes (left) is lost in inception editants (right) as quantified by analysis of mean grey values per eye. Boxplot shows median with boundaries representing the 25th and 75th percentiles. Whiskers extend to maximum 1.5 times interquartile range. (C) Illumina amplicon sequencing reveals high abundance of expected edits. (D) Splice-acceptor removal (CAG>CAA, black outlined box) in *rx3* via intronic *de novo* PAM generation (ABE8e, GAG>GGG) and subsequent C-to-T editing (evoBE4max, sequence adjusted (red outlined box) *rx3-inception-adjusted* guide RNA). (E) At stage 26 eye development is apparent in wild-type (WT) embryos (left), but dramatically affected in *rx3-*inception editants (right, 37.5 %). (F) Illumina amplicon sequencing revealed high rates of the anticipated splice-acceptor mutation. (G) Local multi-codon editing via inception (both non-synonymous) at the *kcnh6a* locus to create hypomorphic alleles with ABE8e only and two guide RNAs (sequence adjusted (red outlined box) *kcnh6a-inception-adjusted*). (H) At 4 dpf the two-chambered embryonic WT heart displays strong Diastole/Systole periodicity (regularly patterned kymograph, used line spanning ventricle (v) and atrium (a) indicated in left panel). *kcnh6a-*inception editants displayed heart phenotypes (26.9 %) including heart morphology, reduced ventricular contractility and the exemplary displayed 2:1 atrioventricular block (white brackets in kymograph (line indicated in left panel), Movie S1). (I) Illumina amplicon sequencing revealed high abundance of expected nucleotide changes. AA pos, amino acid position; canonical PAM (orange); *de novo* PAM (green); inception efficiency, ratio of highest nucleotide changes (step 2 / step 1 * 100); nt, nucleotides; red box, sequence adjustment in inception guide RNAs

Upon injection of the *oca2*-inception mix into one-cell stage embryos efficient loss-of-RPE-pigmentation was detected at 4.5 days post fertilization (dpf) (Fig. 2B; Table S2). PCR amplification and Illumina sequencing of the targeted *oca2* locus revealed efficient *de novo* PAM generation (c.1011A>G) in 79.7 % and the incepted PTC introduction (Gln333*) in 26.1 % (Fig. S1, Table S3). As intended, changes in the canonical base editing window predominantly led to synonymous changes establishing a novel PAM (63.6 % Ala337Ala) and subsequent non-synonymous edits at the initially inaccessible target-site (41.7 % Thr332Ile, 26.1 % Gln333*). Strikingly, although the inception mix contained adenine and cytosine base editors that could edit at both sites, they showed a higher activity at the respective intended target-site (Fig. 2C; Fig. S1, Table S3). To test whether the guide RNAs would cause loss-of-pigmentation in the RPE independently, control injections with both editors and either one of the two guide RNAs were performed and resulted in almost wild-type pigmentation (Table S2, Fig. S2). Illumina sequencing of the *oca2-newPAM* control injection revealed 85.1 % *de novo* PAM generation (c.1011A>G, Fig. S1) and rare indel events (1.2 % of reads; Fig. S2). The *oca2-inception* control injection underscored that in the absence of a proper PAM, binding of the editor to the DNA fails: Thr332 and Gln333 codons remained unaltered in 93.6 % and 96.5 % of cases (Fig. S2, Table S3).

Next, we generated splice-site mutations as an efficient alternative to establish knock-outs (Garcia-Tunon et al., 2019). We used inception to target the splice-acceptor-site of coding exon 2 of the *retinal homeobox transcription factor 3* gene (*rx3*) known to cause eyeless phenotypes (Loosli et al., 2001, Zilova et al., 2021) (Fig. 2D). In 38 % of the injected *rx3-*inception editants, eyes were lost or dramatically underdeveloped (Fig. 2E; Table S2) as a result of efficient *de novo* PAM generation (42 %) and subsequent mutation of the CAG to CAA splice-acceptor-site (38 %, Fig. 2F) while in the single guide controls, impaired eye formation was only detected in rare cases (*rx3-newPAM* mix, 2 %; *rx3-inception* mix, 4 %; Fig. S2, Table S2).

Hypomorphs are often employed to overcome the lethality of null mutants in developmental or cellular key genes (Peterson and Murray, 2022). Thus, we targeted *kcnh6a* (potassium voltage-gated channel, subfamily H (eag-related), member 6a), a key gene controlling heart contraction, in its highly conserved and mutation sensitive membrane spanning S4 domain (Cornean et al., 2022, Hoshijima et al., 2019). Introduction of multiple non-synonymous codon substitutions should thus correlate with severity of detectable phenotypes. To accumulate locally confined multi-codon edits we designed a pair of guide RNAs to cause LysThr506-507ArgAla and Ile502Val by inception editing (Fig. 2G). Changes introduced in the canonical base editing window required sequence adjustment for the *kcnh6a-inception* guide RNA to bind and facilitate editing in the second step (c.1519A>G, position 1 of the *kcnh6a-inception* target-site; Fig. S3). Injection of the kcnh6a-inception mix containing the adjusted inception guide RNA revealed 27 % of heart phenotypes (Table S2) comprising 2:1 atrioventricular block and reduced ventricular contractility similar to earlier reports (Cornean et al., 2022) (Fig. 2H, Movie S1). Illumina sequencing of phenotypic editants confirmed efficient *de novo* PAM generation (33.2 %, c.1519A>G) as well as the intended codon changes (Lys506Arg, 3.5 %; Thr507Ala, 80.6 %; Ile502Val, 28.0 %; Fig. 2I and Table S3). Injection of the control mixes only caused phenotypes in rare cases (Fig. S2). The *kcnh6a-inception* control was literally devoid of mutations due to the absence of the canonical editing event (Fig. S2). Although *de novo* PAM mutation was efficiently introduced (Thr507Ala, 81.5 %) in the *kcnh6a-newPAM* control phenotypes were rare (Fig. S2). Taken together this highlights the phenotypic boost by inception-mediated multiple codon changes.

Overall, inception showed efficient sequential editing events in a one-shot approach at all targeted loci. The rate-limiting step for inception always was the *de novo* PAM generation. Once established, the second edit occurred at almost quantitative rates (step 2 / step 1 * 100; *oca2*, 75.5 %; *rx3*, 91.0 %; *kcnh6a*, 88.6 %; Fig. 2). In essence we show that using base editors in a sequential manner already widens the scope of base editing using the established tools with NGG PAM recognition. Any first editing event introducing a new PAM-site presents a substrate for inception. This opens unlimited possibilities regarding combinations of different PAMs and base editors broadening the targeting range without compromising target specificity.

## Materials and Methods

### Fish maintenance

Adult medaka fish (*Oryzias latipes*, Cab strain) were bred and maintained as closed stocks at 28°C on a 14h:10h light:dark cycle at the Heidelberg University. Fish husbandry and experiments were completely in accordance with the local animal welfare standards (Tierschutzgesetz §11, Abs. 1, Nr. 1, husbandry permit number 35-9185.64/BH Wittbrodt).

### Base editor plasmids and mRNA synthesis

Following plasmids were used in this study: pCS2+_evoBE4max (Cornean et al., 2022), pCMV_AncBE4max (Addgene plasmid #112094) and pCMV_ABE8e (Addgene plasmid #138489) were gifts from David Liu.

ABE8e and ancBE4max plasmids were linearized using SapI (NEB), the evoBE4max plasmid was linearized with NotI-HF (NEB). The digests were purified with QIAquick PCR Purification Kit (Qiagen). *In vitro* transcription of mRNAs was performed using the mMESSAGE mMACHINE SP6 or T7 Transcription Kit (Invitrogen) and purified with the RNeasy Mini Kit (Qiagen), according to manufacturers’ protocols. The quality of the mRNA was assessed with an RNA test gel.

### sgRNAs and crRNAs

All guide RNAs (*oca2, rx3, kcnh6a*) were checked for off-targets using CCTop (Stemmer et al., 2015) and ACEofBASES (Cornean et al., 2022) with standard parameters (Stemmer et al., 2015, Cornean et al., 2022). Cloning of single guide RNA (sgRNA) templates and transcription was performed as described (Stemmer et al., 2015). Target specific crRNAs and tracrRNAs were ordered from IDT (custom Alt-R crRNA). crRNA (100 µM) and tracrRNA (100 µM) were diluted in nuclease-free duplex buffer (IDT) to a final concentration of 40 µM and incubated at 95°C for 5 min.

Following guide RNAs were used in this work (in 5’-3’ direction, PAM in brackets): *oca2-newPAM* sgRNA, TTGCAGGAATCATTCTGTGT[GGG] (Hammouda et al., 2019); *oca2-inception* sgRNA, GGAAACCCAGGTGGCCATTG[CAG]; *kcnh6a-newPAM* sgRNA, TGAAGACAGCCCGACTGCTC[AGG] (Cornean et al., 2022); *kcnh6a-inception-original* crRNA, TCTCATTGGATTACTGAAGg[CAG]; *kcnh6a-inception-adjusted* crRNA TCTCATTGGATTACTGAAGg[CAG]; *rx3-newPAM* sgRNA, AGCAGAGCGCGCAAAGAACC[AGG] (Zilova et al., 2021); *rx3-inception-adjusted* crRNA, CCGGCGCTGTGAGGAGAGCg[GAG].

### Microinjections

Microinjections were performed in wild-type Cab embryos at the one-cell stage. Fertilized embryos were injected with an injection mix (Table S1). After injections, embryos were kept in medaka embryo rearing medium (ERM, 17 mM NaCl; 40 mM KCl; 0.27 mM CaCl2•2H2O; 0.66 mM MgSO4•7H2O; 17 mM HEPES) and incubated at 28°C or at 18°C for *rx3*-targeted editants. Embryos were screened for GFP expression 6 hours or 1 day after injection on a Nikon SMZ18 stereomicroscope. Only GFP positive and properly developing embryos were continued with (Table S2).

### Image acquisition and phenotyping

For analysis of *oca2* knock-outs, the embryos were fixed 4.5 days post fertilization (dpf) (Iwamatsu, 2004) in 4 % paraformaldehyde in 1x PBS (137 mM NaCl; 2.7 mM KCl; 240 mg/l KH2PO4; 1.44 g/l Na2HPO4). Images of eyes were acquired with the ACQUIFER Imaging Machine (DITABIS AG, Pforzheim, Germany) and the mean grey value per eye was quantified as previously described (Thumberger et al., 2022). Embryos injected with guide RNAs targeting *rx3* and *kcnh6a* were imaged 4 dpf or 9 dpf with a Nikon digital DS-Ri1 camera mounted onto a Nikon Microscope SMZ18 equipped with the Nikon Software NIS-Elements F version 4.0.

### Genotyping via Sanger sequencing or targeted amplicon sequencing by Illumina

For genotyping, up to eight embryos were ground and lysed in DNA extraction buffer (0.4 M Tris/HCl pH 8.0; 0.15 M NaCl; 0.1% SDS; 5 mM EDTA; pH 8.0; 1 mg/ml proteinase K) at 60°C overnight. Samples were diluted 1:2 with nuclease-free water and proteinase K was heat-inactivated at 95°C for 20 min.

Genotyping samples of *kcnh6a* editants submitted to Sanger sequencing (Eurofins Genomics) were PCR amplified using Q5 High-Fidelity DNA Polymerase (New England Biolabs) and locus-specific primers (kcnh6a_fwd 5’-GCTTTGCAAGGTATAGAGCACAG-3’ and kcnh6a_rev 5’-AACGTTGCCAAAACCCACAC-3’) from pools of eight embryos with 30x PCR cycles. PCR-products were gel purified after agarose gel electrophoresis with Monarch DNA Gel Extraction Kit (New England Biolabs) and submitted for sequencing. The results were analyzed with EditR (1.0.10).

Samples genotyped by Illumina based amplicon sequencing were prepared by pooling multiple amplicons into a single reaction. For *oca2*-inception injections two replicates with eight randomly picked editants were genotyped, for the *oca2-newPAM* control injection one pool of eight randomly picked embryos were genotyped and for the *oca2-inception* control injection, one single embryo with few pigment-free cells was picked for genotyping.

For *rx3*-inception two pools with four and five phenotypic editants were genotyped, for the *rx3-newPAM* and *rx3-inception* control injection one pool of five randomly picked embryos were genotyped for each injection mix.

For *kcnh6a* inception injections two pools with five phenotypic editants were genotyped, for the *kcnh6a-newPAM* and *kcnh6a-inception* control injection one pool of five randomly picked embryos were genotyped for each injection mix.

The three partial loci *oca2, rx3* and *kcnh6a* were PCR amplified with Q5 polymerase (New England Biolabs) and locus-specific primers 5’ extended with partial Illumina adapter sequences (see table below). PCR products were extracted with the Monarch DNA Gel Extraction Kit (New England Biolabs) after running on an agarose gel. PCR products from each locus were pooled to equimolarity at 20 ng/µl and submitted to GeneWiz (Azenta Life Sciences) for sequencing (Amplicon-EZ: Illumina MiSeq, 2 × 250 bp sequencing, paired-end). Sequencing data was analyzed using CRISPResso2 v.2.1.2 (Clement et al., 2019) and its CRISPRessoPooled tool in the Amplicon Mode.

Primers with partial Illumina adapter sequences (underscored):

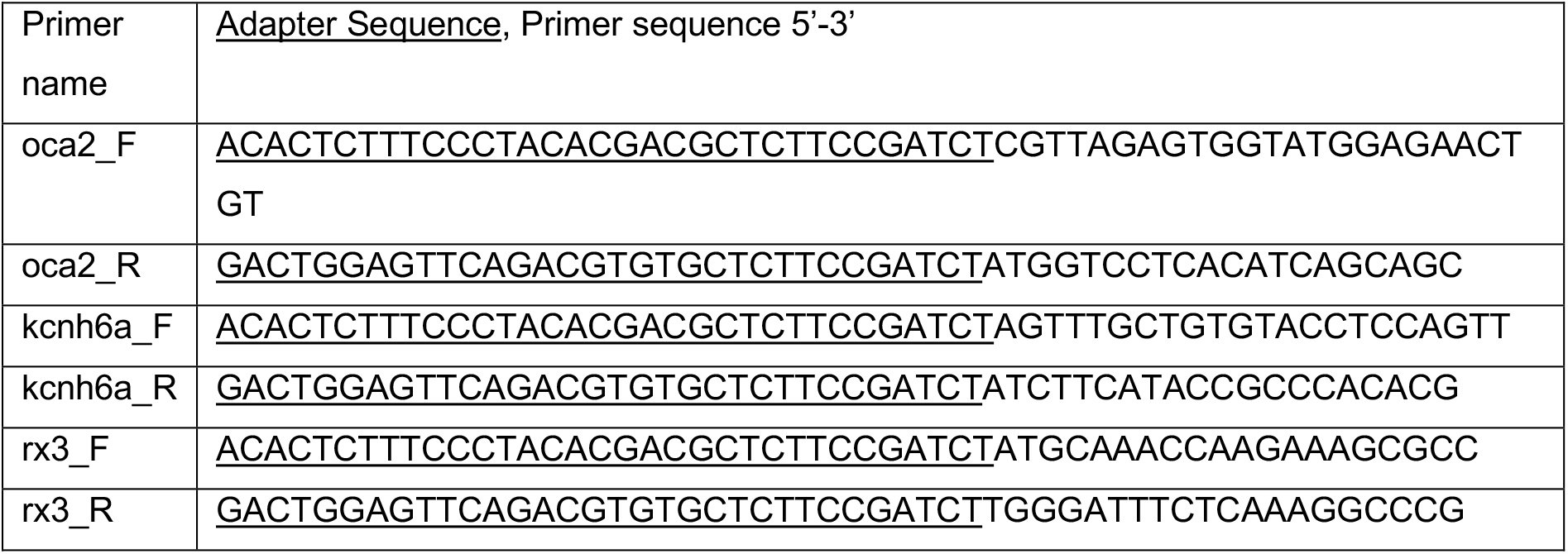

## Data visualization

Microscopy images were processed using Fiji (Schindelin et al., 2012). Data visualization and analysis were performed with ggplot2 in Rstudio and Geneious Prime (2019.2.3, BioMatters). Figures were assembled in Affinity Designer (1.10.5, Serif (Europe) Ltd (1987)).

## Acknowledgments

We thank T Kellner for sgRNA and base editor mRNA synthesis. We are thankful to M Majewski, E Leist, S Erny and A Saraceno for fish husbandry. We thank S Lemke for extensive figure discussions and all members of the Wittbrodt lab for critical, constructive feedback on the procedure and the manuscript.

## Author contribution

TT conceived the inception concept and planned the experimental outline together with KP and JW. TT and KP performed all injection experiments. KP acquired all data with contributions by TT. KP performed all genotyping and sequence analyses. KP and TT performed the formal data analysis and discussed the results with JW. KP, JW and TT wrote the original draft. JW and TT acquired funding sources and TT administered the project.

